# The 13A4 monoclonal antibody to the mouse PROM1 protein recognizes a structural epitope

**DOI:** 10.1101/2022.03.04.482347

**Authors:** Fatimah Matalkah, Scott Rhodes, Visvanathan Ramamurthy, Peter Stoilov

## Abstract

**Purpose:** We endeavored to map the epitope of the rat monoclonal antibody mAB 13A4 to the mouse PROM1 (CD133, AC133) protein. mAB 13A4 is the main reagent used to detect the mouse PROM1 protein. PROM1 is required for the maintenance of primary cilia and mutations in the *Prom1* gene are associated with retinal degeneration.

**Methods:** Epitope tagged clones of splice variants and tiled deletion mutants were used to map the mAB 13A4 epitope and test the predicted tertiary structure of PROM1. The proteins were expressed in Neuro 2a cells and analyzed by Western blot with antibodies to PROM1 and the epitope tag.

**Results:** Deletions in the second and third extracellular domains of the PROM1 protein disrupted the mAB 13A4 epitope. Furthermore, the affinity of mAB 13A4 to the major PROM1 isoform in photoreceptor cells is significantly reduced due to the inclusion of a photoreceptor-specific alternative exon in the third extracellular domain. Interestingly, a deletion in the photoreceptor specific isoform of six amino acids adjacent to the alternative exon restored the affinity of mAB 13A4 to PROM1.

**Conclusion:** mAB 13A4 recognizes a structural epitope that is stabilized by two of the extracellular domains of PROM1. The results of our mutagenesis are consistent with the computationally predicted helical bundle structure of PROM1 and point to the utility of mAB 13A4 for evaluating the effect of mutations on the PROM1 structure. We show that the PROM1 isoform composition needs to be considered when interpreting tissue and developmental expression data produced by mAB 13A4.

## INTRODUCTION

Prominin (PROM1, CD133, AC133) was identified as the antigen of monoclonal antibodies raised against human hematopoietic progenitors and mouse neuroepithelial cells ^1–3^ PROM1 is a glycosylated membrane protein with five transmembrane and three extracellular domains ^1,2^. PROM1 is a member of a conserved family of proteins involved in modulating the architecture of cellular protrusions, such as microvilli and cilia ^4–6^. Mutations in the human Prom1 gene have been reported in various types of retinal degeneration and are primarily associated with cone-rod dystrophy ^7–10^. Mouse models lacking *Prom1* or expressing the dominant Arg373Cys mutant recapitulate the retinal degeneration phenotype and display defects in disk morphogenesis ^9,11^. Most mutations in *Prom1* are recessive and result in loss of function due to premature stop codons. Notably, three missense mutations, Leu245Pro, Arg373Cys, and Asp829Asn, have dominant inheritance patterns ^7^.

The *Prom1* gene produces multiple splicing isoforms that can be tissue, and cell type specific ^12–14^ Six alternative exons in the mouse *Prom1* can potentially produce 24 splice variants, although to date only eight have been enumerated ^12,13^. In mouse photoreceptor cells a microexon, exon 19a, introduces 6 amino acids in the photoreceptor-specific SV8 isoform of *Prom1* ^13,14^. While this exon is present in most vertebrate clades, it is not used in the primates, including humans, due to mutations that disrupt either the 3’ or the 5’ splice site of the exon (Figure 2B).

**Figure 1:**
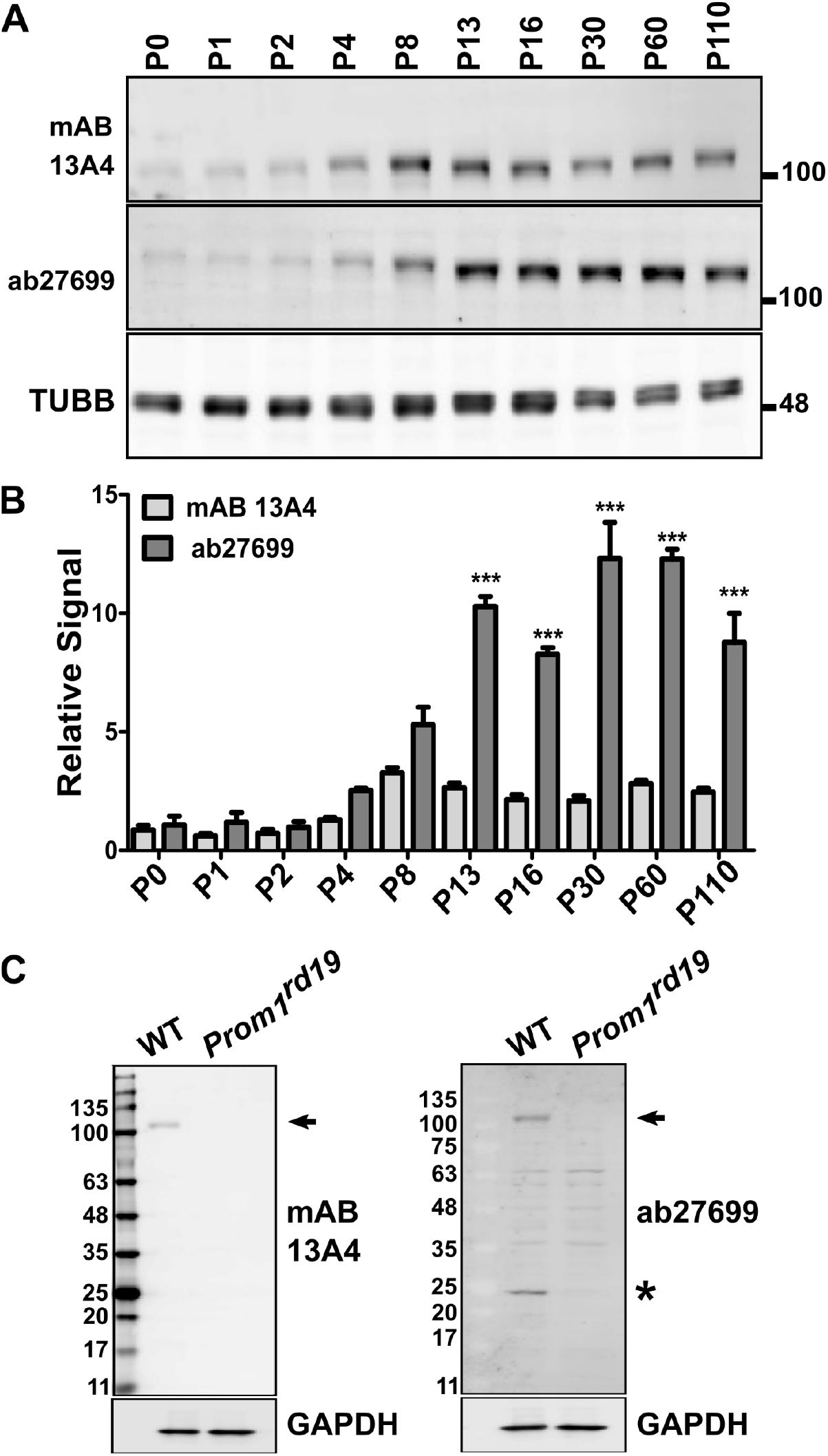
Discrepancy in the levels of PROM1 as determined by the mAB 13A4 ab27699 antibodies. **A)** Immunoblotting for PROM1 in mouse retina lysate collected between postnatal day 0 (P0) and postnatal day 110 (P110) using the mAB 13A4 and the ab27699 antibodies. TUBB serves as a loading control. **B)** Quantification of Prom1 level in retina lysates using the mAB 13A4 and the ab27699 antibodies. Error bars represent the standard error of the mean (n=3). Two-way ANOVA was used to assess the effect of the postnatal day and the antibody used on the PROM1 signal. The statistical significance of the signal mAB 13A4 compared to ab27699 for each day was calculated by Tukey HSD post-hoc test. Tukey HSD p-values of less than 0.001 from the post-hoc test are indicated by “***”. **C)** Test of the specificity of mAB 13A4 and ab27699 antibodies for detecting PROM1 using the retinal lysate from wild type mice and *Prom1^rd19^* mutants that do not express PROM1 protein. Arrow indicates the position of the expected PROM1 band. A 25 KDa band, labeled by asterisk, is detected in the wild type but not *Prom1^rd19^* retina by ab27699. GAPDH serves as a loading control.

**Figure 2:**
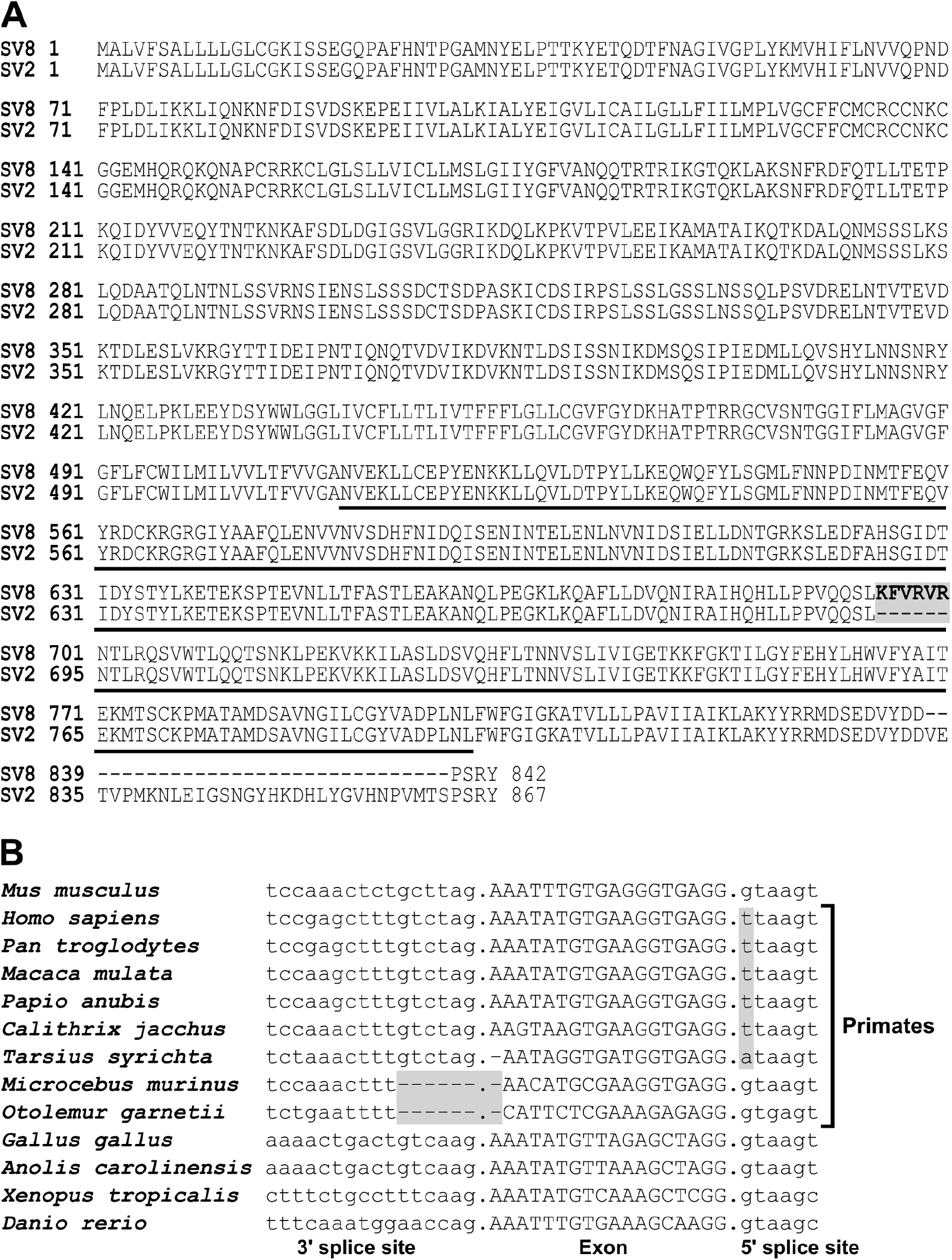
Photoreceptor specific splice variant of PROM1. **A)** Alignment of the photoreceptor-specific SV8 isoform of PROM1 (RefSeq NP_001157057) to the ubiquitously expressed isoform SV2 (RefSeq NP_001157049). The amino acids encoded by the photoreceptor specific exon 19a are shown in bold and shaded in gray. The third extracellular domain is underlined. **B)** Alignment of vertebrate exon 19a sequences including the adjacent 3’ and 5’ splice sites. Mutations inactivating the splice sites in primates are shaded in gray.

Despite playing conserved and critical functions the Prominin genes show relatively low conservation of their primary amino acid sequence. For example, human and mouse PROM1 proteins share approximately 61% sequence identity. Consequently, antibodies to PROM1 tend to be species specific. In mice, the rat monoclonal antibody mAB 13A4 is the reagent typically used to detect PROM1. The mAB 13A4 antibody was raised against extract from mouse neuroepithelium ^2^. Its antigen was cloned by screening for reactivity with mAb 13A4 of a phage library of mouse kidney cDNA ^2^. mAB 13A4 is speculated to recognize part of the third extracellular domain of PROM1 because of a truncating mutation in that domain that abolishes its binding, but the epitope was never mapped ^8,15^.

Prompted by a discrepancy between the PROM1 protein levels measured in the postnatal retina by mAB 13A4 and the mouse monoclonal antibody to PROM1 ab27699, we set out to map the epitope of mAB 13A4. We found that mAB 13A4 recognizes a structural epitope that can be disrupted by deletions in the second and third extracellular domains. Furthermore, the affinity of mAB 13A4 to PROM1 is dramatically reduced by the inclusion of the photoreceptor-specific exon 19a.

## MATERIALS AND METHODS

### Animals

The *Prom1^rd19^* mice were acquired from the Jackson laboratory (B6. BXD83-Prom1^rd19^/Boc, Stock No: 026803). All experiments were conducted with the approval of the Institutional Animal Care and Use Committee at West Virginia University.

### *Prom1* clones

We obtained a full length Mammalian Gene Collection clone of the mouse photoreceptor specific isoform (SV8) of *Prom1* from Horizon Discovery (clone ID: 4502359, NCBI accession: BC028286). Gibson assembly (NEB# E5510) was used to generate Flag-tagged *Prom1* clones and deletion mutants in pcDNA3.1+ (Invitrogen). The primers used for cloning are listed in Supplementary Table 1.

### Cell culture and transfection

Mouse Neuro-2a (N2a) neuroblastoma cells (ATCC CCL-131) were cultured in OptiMEM reduced serum media buffered with sodium bicarbonate and supplemented with 4% (v/v) fetal bovine serum (FBS, R&D Systems, Minneapolis, MN). The cells were grown at 37 °C in a 5% CO2 humidified atmosphere. The cDNA clones were transiently transfected in N2a cells using polyethylenimine ^16^. Cell lysates for Western blot analysis were collected at 24 hours post-transfection.

### Denaturing Gel Electrophoresis and Western Blot

Flash-frozen mouse retinas and N2a cells transiently transfected with Prom1-expressing constructs were lysed in RIPA buffer (50 mM Tris HCl-PH 8.0, 150 mM NaCl, 1.0% TritonX-100, 0.5% Sodium Deoxycholate, 0.1% SDS) supplied with protease (Sigma-Aldrich catalog# 535140-1ML) and phosphatase inhibitors cocktail (Sigma-Aldrich catalog # P5726-1 ML). After homogenization, the lysate was incubated on ice for 10 mins, then cleared by centrifugation for 15 mins. 20 μg of protein extract was resolved in 4-20% polyacrylamide SDS–PAGE gel and transferred onto polyvinylidene difluoride (PVDF) membranes (Immunobilon-FL, Millipore). After blocking with BSA in PBST (Phosphate-buffered saline with 0.1% Tween-20), the membranes were probed with primary antibodies overnight at 4°C, followed by incubation with fluorescently labeled (Alexa Fluor 647 or 488, Jackson ImmnuoResearch) secondary antibodies for 1 hour at room temperature. The membranes were then imaged on Amersham Typhoon Phosphorimager (GE Healthcare).

### Blue Native Polyacrylamide Gel Electrophoresis (BN PAGE)

The samples were lysed in BN PAGE sample buffer (Thermo Fisher Catalog# BN2008) containing 1% digitonin and protease inhibitors following the manufacturer’s recommendation. The lysates were treated with benzonase at room temperature for 30 minutes to shear the DNA, and cleared by centrifugation for 15 mins. Prior to electrophoresis, the samples were mixed with Coomassie G-250 and resolved in 3-12% NativePAGE Bis-Tris gel (Invitrogen Catalog #BN1003BOX) as per the manufacturer’s recommendations. The gels were then transferred on PVDF membranes (Immunobilon-FL, Millipore) following the manufacturer’s recommendations. After transfer, the membranes were incubated in 8% acetic acid for 15 minutes to fix the proteins, rinsed with deionized water, and air-dried. The membranes were blocked, probed with antibodies, and imaged as described above in the denaturing gel electrophoresis protocol.

### Antibodies

The primary and secondary antibodies that were used throughout our studies include the following: mouse anti-β-tubulin (1:5000; catalog # T8328-200ul, Sigma-Aldrich, St. Louis, MO), mouse anti-GAPDH (1:20,000; custom made), mouse anti-flag M2 (1:1000, catalog # F1804-200UG, Sigma-Aldrich, St. Louis, MO), rat anti-Prom1(1:1000, clone ID:13A4, ThermoFisher, Waltham, MA), mouse anti-GFP HRP conjugated GFP tag ( 1:1000, Cat #HRP-66002, ThermoFisher, Waltham, MA), mouse anti-Prom1 (1:1000, Cat #ab27699, Abcam, Cambridge, MA), Alexa Fluor 647 conjugated AffiniPure Goat-Anti rabbit IgG (1:3000, Jackson ImmunoReserach, West Grove, PA), Alexa Flour 488 conjugated AffiniPure Goat-Anti mouse IgG (1:3000, Jackson ImmunoReserach, West Grove, PA), and Alexa Fluor 647 conjugated AffiniPure Goat-Anti rat IgG (1:3000, Jackson ImmunoReserach, West Grove, PA).

### Statistical analysis

The two-way analysis of variance (ANOVA) with Tukey Honest Significant Differences (HSD)post-hoc test or two-tailed unpaired Student’s T-test were used to determine statistical significance as indicated in the results section. Quantitative data is presented as the mean of three biological replicates ±standard error of the mean.

#### Protein structure prediction and visualization

The RobeTTa structure prediction service was used to create models of the photoreceptor-specific SV8 (RefSeq NP_001157057) isoform that contains exon 19a and the ubiquitously expressed isoform SV2 (RefSeq NP_001157049) that lacks exon 19a (see Figure 2A for alignment of the two sequences). To create images of the structures we used Pymol (The PyMOL Molecular Graphics System, Version 2.0 Schrödinger, LLC).

## RESULTS

### Discrepancy in the PROM1 protein levels measured by the mAB 13A4 and ab27699 antibodies

While investigating the expression of PROM1 in postnatal mouse retina we noticed that when measured by mAB 13A4 the PROM1 protein levels peaked at postnatal day 8, five days before the peak recorded by ab27699 (Figure 1A and B). Furthermore, mAB 13A4 showed approximately three to five fold lower levels of PROM1 at postnatal days 13 and beyond compared to ab27699 (Figure 1A and B). Two way ANOVA showed significant effect of the day after birth (F(9)=55.26, p-value<2*10^-16^) and the antibody used (F(1)=365.48, p-value<2*10^-16^) on the measured PROM1 protein levels. According to the Tukey HSD post-hoc test the signals detected by mAB 13A4 and ab27699 were significantly different starting from postnatal day 13. It is possible that cross-reactivity of ab27699 to other proteins of size similar to PROM1 in the retina could have compromised its performance. To rule out cross-reactivity we probed retinal extracts from wild type and homozygous *Prom1^rd19^* mutant retina with mAB13A4 and ab27699. The *Prom1^rd19^* allele contains a premature stop codon that abolishes the expression of the PROM1 protein ^17^. Both antibodies recognized a protein just over 100KDa in size in the wild type retina that was not detected in the extract from *Prom1* knockout retina (Figure 1C). Thus, the discrepancy in the signal between mAB13A4 and ab27699 is likely due to differences in the availability of the epitopes recognized by the two antibodies.

### Reduced affinity of mAB13A4 to PROM1 carrying the photoreceptor-specific exon 19a

A short 18nt microexon, exon 19a, is included in the *Prom1* transcripts specifically in photoreceptor cells (Figure 2) ^14^. The inclusion rate of exon 19a starts to increase at postnatal day 3 and the exon 19a containing transcripts become dominant in the retina after postnatal day 8 ^14^. The peptide encoded by exon 19a is inserted in the third extracellular loop of PROM1 (Figure 3A), which is also the proposed location of the mAB 13A4 epitope ^8,15^. Thus it was possible that inclusion of exon 19a disrupts the epitope of mAB 13A4 while leaving intact the epitope of ab27699. To determine if exon 19a disrupts the mAB 13A4 epitope we generated Flag-tagged cDNA clones that either contain (SV8) or skip (SV8(-Ex19a)) the photoreceptor specific exon 19a. The cDNA were transfected in N2a cells and their expression was probed with mAB 13A4, ab27699 and anti-Flag antibodies. As expected, mAB 13A4 showed significant reduction in its affinity to the cDNA containing exon 19a compared to the epitope tag (Figure 3B). The inclusion of exon 19a had no effect on the affinity of the ab27699.

**Figure 3:**
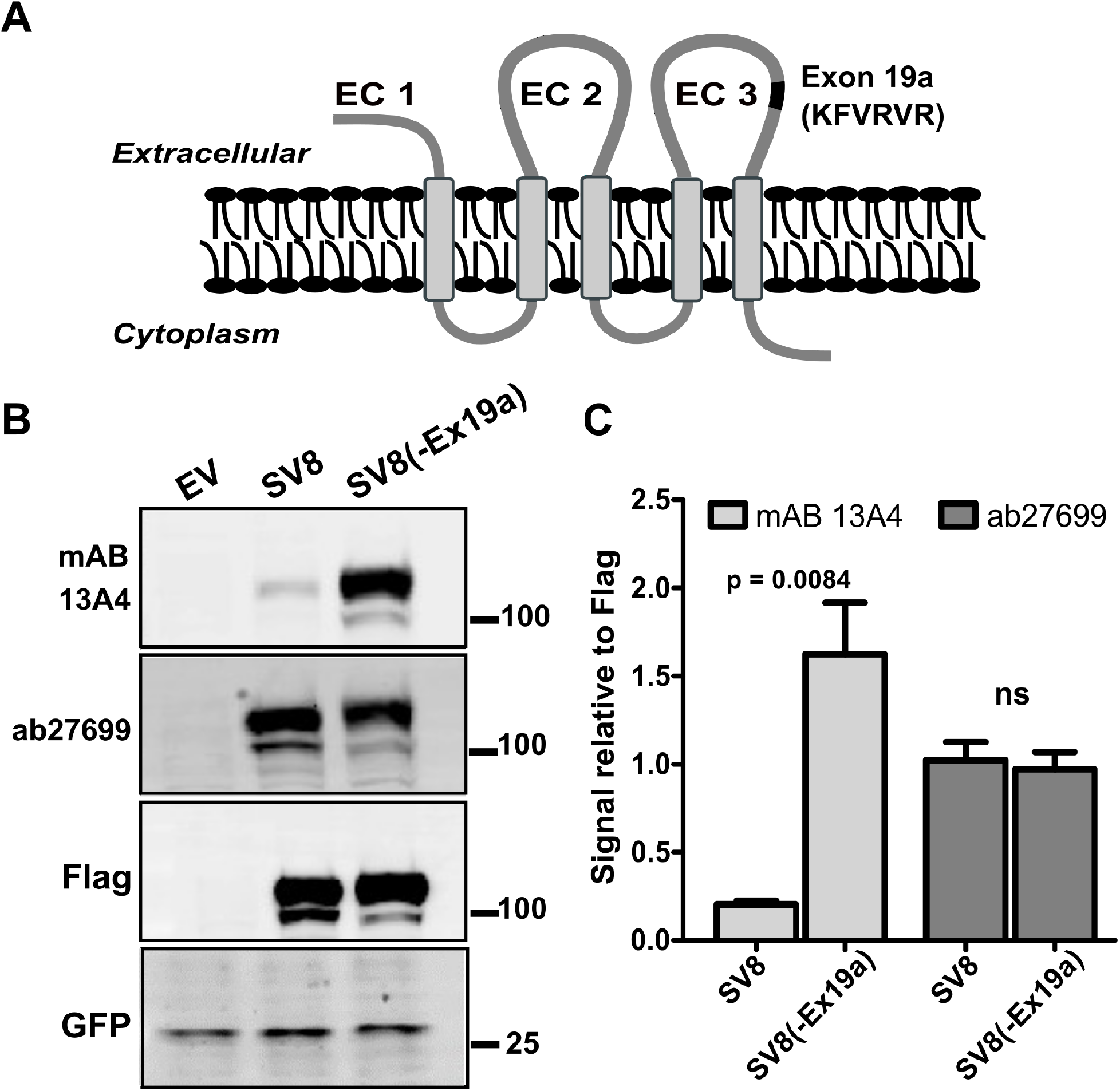
The affinity of mAB 13A4 to PROM1 is affected by alternative splicing. **A)** Schematic representation of the PROM1 structure showing the position of exon 19a (black) in the third extracellular domain (EC 3) of the photoreceptor specific PROM1 isoform SV8. Extracellular domains one through three are labeled as EC 1, EC 2 and EC 3 respectively. **B)** Western blot of recombinant SV8 and SV8 lacking exon 19a, SV8(-Ex19a), expressed in N2a cells with mAB 13A4, ab27699 and antibody to the Flag epitope. Transfection with empty pcDNA3.1 vector (EV) was used as a negative control. All transfections were spiked with vector expressing GFP to control for transfection efficiency. **C)** Quantification of mAB 13A4 and ab27699 signals relative to the signal from the Flag-tag antibody. Error bars represent the standard error of the mean (SEM, n=3). Unpaired Student’s t-test was used to assess the statistical significance and the p-value is indicated on the chart.

### mAB 13A4 recognizes a structural epitope

To map the epitope of mAB13A4 we created a series of tiled deletion mutants of the SV8(-Ex19a) cDNA, that originated at the point where exon 19a would have been inserted and progressed in C-terminal and N-terminal direction (Figure 4A). The deletion mutants covered 108 amino acids of sequence. Nine out of ten deletions resulted in complete loss of the mAB 13A4 epitope (Figure 4B). Only the most C-terminal deletion in the series (D+4) could be detected by mAB 13A4. The results of the deletion mutagenesis indicate an epitope for mAB 13A4 that is at least 94 amino acids in length. This length far exceeds the size range of linear peptide epitopes and strongly argues for a structural rather than a linear epitope ^18,19^.

**Figure 4:**
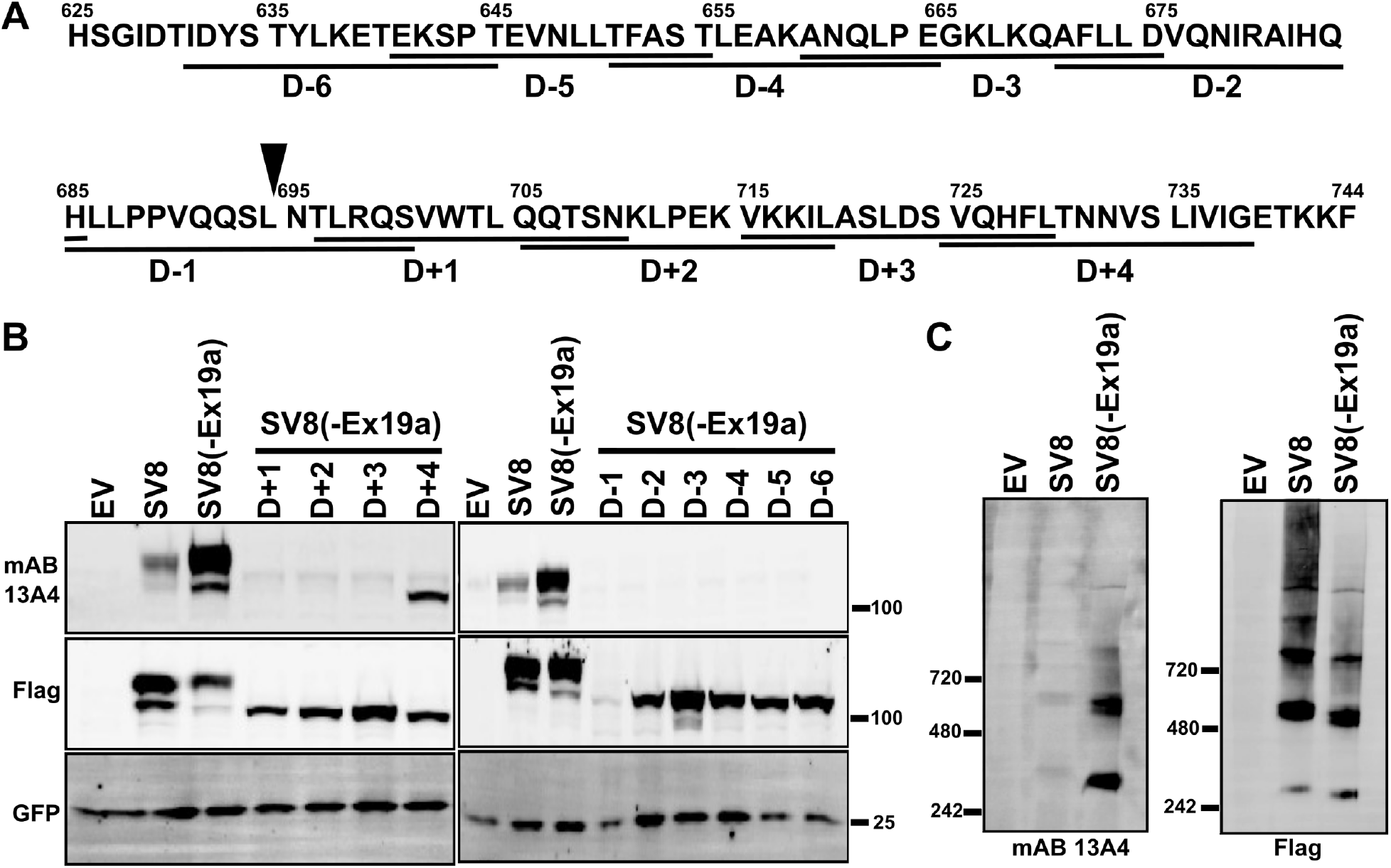
Mapping of the mAB 13A4 epitope. **A)** Sequence of the third extracellular domain. Underlining shows the positions of the deletions used for epitope mapping. The deletion variants were generated starting from the SV8 clone lacking exon 19a, SV8(-Ex19a). Solid triangle above the sequence shows the position where exon 19a is inserted in the photoreceptor specific isoform. **B)** Western blot analysis of PROM1 deletion mutants expressed in N2a cells using mAB 13A4 and Flag-tag antibodies. All mutants with exception of D+4 resulted in loss of the mAB 13A4 epitope. Transfection with empty pcDNA3.1 vector (EV) was used as a negative control. All transfections were spiked with a vector expressing GFP to control for transfection efficiency. **C)** Native gel Western blot analysis using mAB 13A4 and Flag-tag antibodies of N2a cell lysate transfected with either an empty vector (EV), SV8, or SV8(-Ex19a).

The western blots shown on Figures 1, 3, and 4 were performed using denaturing gel electrophoresis. Detecting a structural antigen using this approach would require the protein to renature on the membrane prior to probing with the primary antibody. Consequently, our results may reflect the propensity of the splicing isoforms and deletion mutants to renature rather than the genuine affinity of the mAB 13A4 to the native proteins. To test if this is the case we resolved the proteins produced by the SV8 and SV8(-Ex19a) clones on native gel electrophoresis and probed the membranes with mAB 13A4 and anti-Flag antibodies. The exon 19a containing protein was recognized with significantly reduced affinity (Figure 4C) demonstrating that inclusion of exon 19a alters the structure of PROM1 rather than affecting its folding rate. Interestingly, under native conditions PROM1 formed higher order complexes corresponding in size to dimers and tetramers.

### Computationally derived PROM1 tertiary structure predicts the effect of sequence manipulation on the mAB13A4 epitope

To better understand the nature of the mAB 13A4 epitope and how our deletion mutagenesis affected it we needed a tertiary for PROM1. There are no empirically derived structures of PROM1. Nevertheless, recent advances in computational approaches for structure prediction are producing remarkably accurate structures ^20,21^. We used the RobeTTa structure prediction service to model the structures of the mouse PROM1 isoforms SV8 and SV2 (Figure 2A)^20^. The photoreceptor-specific SV8 isoform differs from canonical SV2 isoforms by the inclusion of exon 19a in the third extracellular domain and the skipping of two exons in the cytoplasmic C-terminal domain. The structures predicted by RobeTTa for the SV8 and SV2 proteins were in excellent agreement with each other and with the structure of the human PROM1 predicted by Alpha Fold ^22^. In the predicted PROM1 structure, the second and third extracellular domain each form two antiparallel alpha helices that are continuous with the adjacent transmembrane domains. The alpha helices formed by the second and third extracellular domains and the adjacent transmembrane domains are packed in a four helix bundle (Figure 5A). Inclusion of exon 19a lengthens the second helix of the third extracellular domain causing a kink in the bundle (Figure 5A). Mapping the positions of the deletions that disrupted the mAB 13A4 epitope on the PROM1 structure showed that they were located towards the middle portion and the tip of the helical bundle. The D+4 mutation, which was the only one that did not result in loss of the mAB 13A4 epitope, was the furthest from the tip of the helical bundle. Based on the positions of the deleted segments and the six amino acids encoded by exon 19a in the PROM1 structure we made three predictions: (i) deleting six amino acids adjacent to exon 19a (Del AA 6) in the photoreceptor-specific PROM1 isoform should shorten the helix compensating the inclusion of exon 19a, and restore the mAB 13A4 epitope (Figure 5B, colored dark blue on the structure of SV8); (ii) Deletion of a 15 amino acid segment (D-7) in the third extracellular domain opposite to D+4, should retain the mAB 13A4 epitope due to its distance from the location of the epitope in the upper half of the helical bundle (Figure 5B, colored red on the structure of SV2); (iii) Deletion of a 15 amino acid segment (Del EC 2) in the upper half of the second extracellular domain of SV8(-Ex19a) should result in loss of the mAB 13A4 epitope (Figure 5B, colored sky blue on the structure of SV2). All three predictions proved to be accurate when the proteins expressed from the corresponding clones were analyzed by Western blot (Figure 5C). Deletion of the six amino acid long segment recovered the mAB 13A4 epitope in the photoreceptor specific SV8 isoform. Conversely, the deletion in the second extracellular domain of the protein encoded by the SV8(-Ex19a) clone resulted in complete loss of the epitope. Finally, deletion D-7 in SV8(-Ex19a) preserved the epitope, although the affinity of mAB 13A4 was reduced. The results of the structure directed mutagenesis provide further support for mAB 13A4 recognizing a structural epitope. In addition, our results demonstrate the utility of the modeled PROM1 structures.

**Figure 5:**
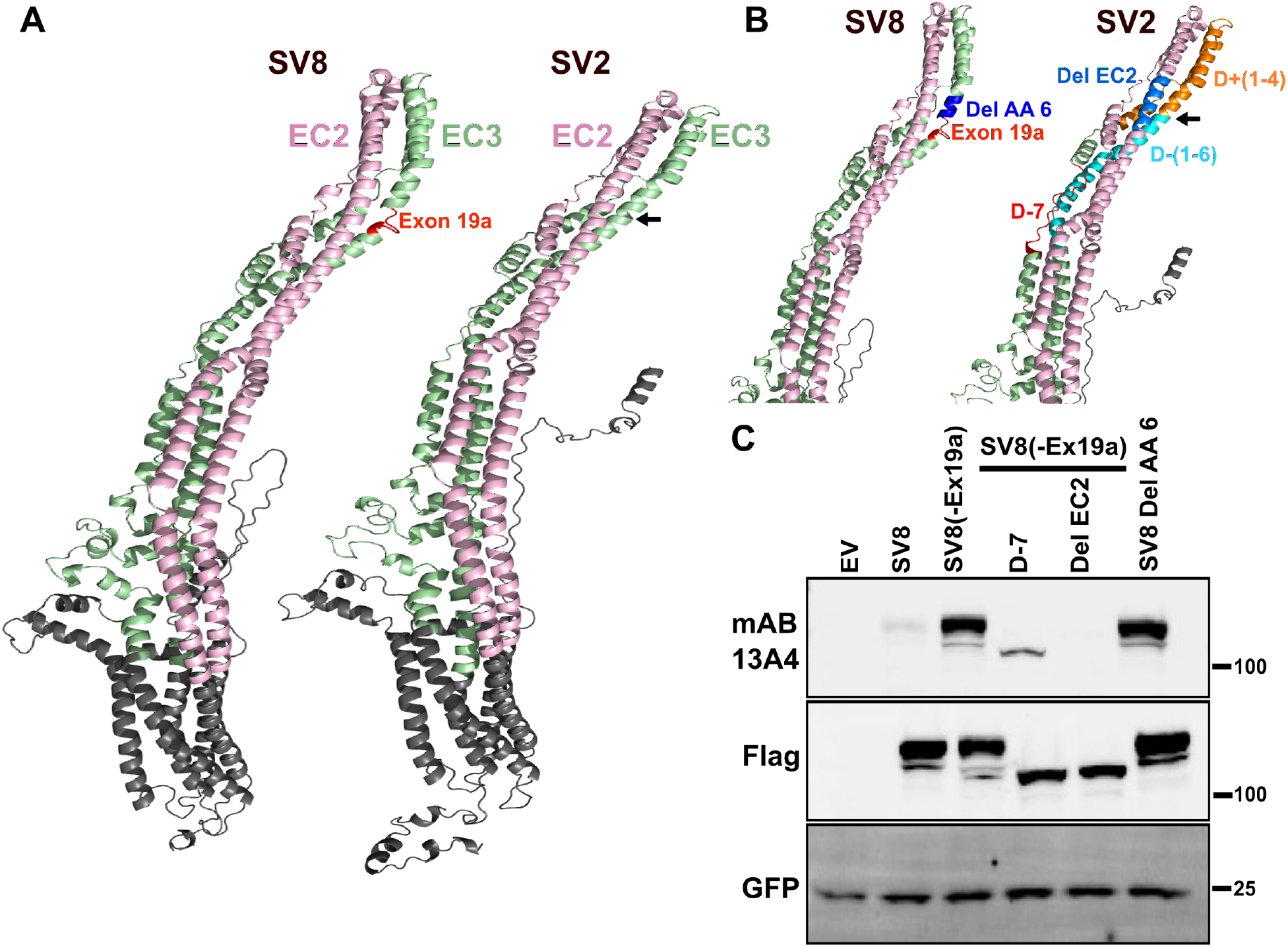
Computationally derived tertiary structure of PROM1 predicts the effect of mutations on the mAB 13A4 epitope. **A)** Structures of PROM1 isoforms SV8 and SV2 showing extracellular domain 2 (EC2) in pink and extracellular domain 3 (EC3) in green. Inclusion of exon 19a in SV8 (red) forms a kink in the helical bundle compared to SV2. Arrow points to the position of the excluded exon 19a in SV2. **B)** Map of the segments deleted in the experiments shown on Figure 4 and on panel C of this figure on the tertiary structure of PROM1. Exon 19a is shown in red on the structure of SV8. The deletion Del AA6 analyzed on panel C is shown in blue on the structure of SV8. On the structure of SV2 cyan color indicates deletions D-1 through D-6 and orange indicates deletions D+1 through D+4 from the experiments shown on Figure 4. Also on the structure of SV2, the positions of the deletions D-7 and EC2 analyzed on panel C are shown in red and sky blue, respectively. Arrow points to the position of the excluded exon 19a in SV2. **C)** Western blot analysis of PROM1 deletion mutants expressed in N2a cells using mAB 13A4 and Flag-tag antibodies. All transfections were spiked with a vector expressing GFP to control for transfection efficiency.

## DISCUSSION

To reliably interpret results from techniques that employ antibodies it is essential to know the antibody epitope, its specificity and its affinity. While mapping the antibody epitope may be important, it is usually not considered necessary as long as specificity to the target can be demonstrated. Such practice leaves a gap that in certain cases can have significant impact on interpreting experimental results as we show here for mAB 13A4. mAB 13A4 is widely used to detect the mouse PROM1 protein because of its excellent specificity and the lack of alternatives with comparable performance. As of the time of writing of this article, there are over 300 publications in Google Scholar citing the 13A4 antibody in the context of PROM1. Here we show that the mAB 13A4 antibody recognizes a structural epitope and its affinity for naturally occuring PROM1 isoforms can vary dramatically. When used to measure PROM1 levels in the retina, mAB 13A4 underestimated the protein amount by a factor of five compared to ab27699.

To determine the exact mAB1 13A4 epitope unequivocally will require determining the structure of the PROM1 - mAB 13A4 complex, which is beyond the scope of the current work. Nevertheless, we show that mAB 13A4 is a useful reagent for detecting perturbation in PROM1 structure as changes to the PROM1 sequence that were hundreds of amino acids apart abolished the mAB 13A4 epitope. Furthermore we created three mutations in PROM1 guided by the computational model of its structure. The effect of these mutations on the mAB 13A4 epitope was in line with the predicted structure, providing empirical evidence for its validity. Finally, we demonstrate that under native conditions PROM1 can form higher order complexes, providing a possible path towards understanding the dominance of Prom1 mutations in patients with cone-rod dystrophy.

## Supporting information

Supplemental Table 1. List of cloning primers

## Acknowledgements

We are grateful to Dr. Maxim Sokolov and Dr. Douglas Kolson for their assistance with native gel electrophoresis.

## Notes

**Funding:** This work was supported by the National Institutes of Health 2R01EY025536, (P.S. and V.M.) and bridge funding provided by the West Virginia University Health, Sciences Center Office of Research and Graduate Education (P.S.).

### Competing Interest Statement

The authors have declared no competing interest.

